# Novel insights into *Entamoeba* cyst wall formation and the importance of actin cytoskeleton

**DOI:** 10.1101/2020.04.24.059360

**Authors:** Deepak Krishnan, Santhoshi Nayak, Sudip Kumar Ghosh

**Author notes:** **Author for correspondence:** Sudip Kumar Ghosh.

## Abstract

The cyst wall of *Entamoeba histolytica,* the causative agent of the amoebiasis, is a potential target for new drugs. The “Wattle and Daub” model of cyst wall formation of *Entamoeba invadens* had already been reported. In this study, we demonstrate in more detail the morphological stages of chitin wall formation in *E. invadens* using fluorescent chitin-binding dyes and immunolocalization of the cyst wall proteins. Here, the expression and localization of chitin synthase and the importance of actin cytoskeleton dynamics at cellular level, during encystations have been demonstrated for the first time. Chitin deposition was found to be initiated on the cell surface mostly from one distinct point, though multipoint initiation was also observed sometimes. From these points, the wall grew outwards and gradually covered the entire cyst surface with time. The initiation of chitin deposition was guided by the localization of chitin synthase 1 on the plasma membrane. The gradual formation of the cyst wall follows the Wattle and daub model. The chitin deposition occurred on the foundation of Jacob lectin at the cell membrane, and the other cyst wall components, like chitinase, and Jessie were also found to be present in the growing incomplete walls. In contrary to the Wattle and daub model, Jessie was found to be expressed and localized in the growing wall at the early hours of encystations. During encystation, F-actin was reorganized into the cortical region within an hour encystation initiation and remained intact until the completion of the chitin wall. Disruption of cortical actin polymerization with 2, 3-Butanedione monoxime inhibited proper wall formation but produced wall-less cysts or cysts with defective chitin wall. Malformations of cyst-walls were mainly due to improper localization and activity of chitin synthases, which indicates the indispensability of cortical actin cytoskeleton for the proper cyst wall formation.

**Author Summary:** *Entamoeba* parasites reach new hosts using the resistant cyst form, so preventing its formation can stop the spread of amoebiasis. The resistant nature of the cyst is due to the chitin wall, and thus identifying the critical steps of wall formation could provide targets for designing new drugs. Here we studied the morphological stages of the cyst wall formation by observing how the chitin and other cell wall components were deposited on the cell surface using fluorescent chitin-binding dyes and antibodies against cyst wall proteins. In most cases, the chitin wall was found to start from one distinct point from which it spread all over the cell surface, guided by chitin synthase. The composition of these incomplete walls was the same as a mature cyst wall indicating that the wall may be a result of extracellular self-assembly of its constituents from one starting point. We have also observed that F-actin polymerized in the cortex of encysting cells and its disruption resulted in wall-less cysts or cysts with aberrant walls showing the importance of actin cytoskeleton in proper chitin deposition.

## Introduction

*Entamoeba histolytica* is the causative agent of amoebiasis, the third most important parasitic disease after malaria and schistosomiasis, and causes 40-50 million cases and 40,000-100,000 deaths annually (1). Nitroimidazole drugs like metronidazole, tinidazole, ornidazole are the most important drugs for amoebiasis, but most of these drugs have serious side effects. Though drug resistance in clinical isolates has not been reported yet, drug-resistant *E. histolytica* have been generated in laboratory condition (2,3) indicating the widespread use of anti-amoebic drugs may eventually cause drug resistance. Thus it is necessary to find new, safer drugs, and for this purpose, novel drug targets are required to be identified. *Entamoeba histolytica* reaches new hosts through the ingestion of water or food contaminated with cysts. Thus developing a chemotherapeutic agent that blocks encystation can prevent the spreading of the parasite by reducing the number of cysts in the environment. The events of chitin cyst wall deposition are of particular interest in this regard as it is the wall that provides resistant nature to the cyst. Immunization with cyst wall proteins of *Giardia lamblia* has been reported to cause significant reduction of cyst shedding in murine models (4–6). To find such drug or vaccine candidates, it is necessary to understand the structure of the chitin wall and the mechanics of its formation.

*E. histolytica* does not encyst in an axenic culture, reptilian parasite *Entamoeba invadens* has been opted as a model to study the encystation as it readily forms cyst when subjected to starvation and osmotic stress (7). *E. invadens* occupy a similar host niche and create similar colon pathology like *E. histolytica* (8). Also, *E. invadens* genome showed considerable sequence similarity to that of *E. histolytica* (9). Most importantly, the cyst characteristics of *E. histolytica* and *E. invadens* are quite similar; both formed a chitin walled tetranucleate cyst with a single chromatoid body. A comparison of gene expression during encystation showed that they share a large number of developmentally regulated genes. Encystation specific genes of *E. invadens* were also found to over-expressed in clinical isolates of *E. histolytica* (10). Immunolocalization experiments have shown that the cyst wall proteins identified in *E. invadens* like chitinase, Jacob lectin, and Jessie are also present in the *E. histolytica* cysts isolated from patients (11).

Unlike the walls of fungi that contain multiple layers and many sugar polymers, the *Entamoeba* cyst wall is single-layered and contains a single sugar polymer, chitin, forming nearly 25% of its dry weight (12). The cyst wall also contains lectins with multiple chitin-binding domains like Jacob, Jessie, and chitinase (13). The assembly of different components into the cyst wall is explained by the “wattle and daub model” (14). According to this model, during the foundation stage, the Jacob lectin is secreted and binds to the constitutively expressed Gal/GalNAc lectins on the cell membrane. During the “wattle” stage, chitin synthases produce the chitin fibrils using UDP-GlcNAc provided by nucleotide sugar transporters (15). The chitin fibrils are then cross-linked by Jacob lectin through their chitin-binding domains (CBD), forming the framework of the wall. Chitinase and chitin deacetylase trim and deacetylate the chitin fibrils (16). Finally, during the daub phase, the Jessie lectin, which has a single CBD and a self-aggregation inducing-C-terminal domain, binds to this framework completing the cyst wall and make it impermeable.

According to the ‘‘wattle and daub’’ model, Jacob lectins are deposited onto the surface in the foundation stage, which then cross-links chitin fibrils in the “wattle stage”, and addition of Jessie lectins occur after 36 hours of encystation in the “daub stage”. However, the expression profiles of cyst wall proteins showed that most of them were over-expressed between 8 and 24 hours (17,18). In order to understand how different components were added to the wall, the intermediate stages of cyst wall formation were investigated. As the *in vitro* encystation is an asynchronous process, and so it is difficult to follow its progression. So we studied the deposition of chitin by staining the cells with chitin-binding dyes and analyzed its maturity based on detergent resistance. Localizing the chitin allowed us to follow the highly asynchronous encystation as chitin is the most important maker of the encystation. We also studied the cyst wall deposition with respect to the appearance of other cyst characteristics like chromatoid body formation and nuclear division.

The relationship between the actin cytoskeleton and chitin wall formation is well established in fungi. Actin was found at sites where the cell wall synthesis is active like the bud, growing apices, and sites of septum formation (19). In regenerating protoplasts of the *Saccharomyces cerevisiae,* actin patches were found over the surface on which a new cell wall was being synthesized (20). During encystation, *E. invadens* undergo morphological changes from highly motile trophozoites to an immobile spherical cyst, which shows the involvement of cytoskeleton in the stage conversion. In the case of *Entamoeba,* morphology and motility are entirely controlled by the actin cytoskeleton as cytoplasmic microtubules are absent (21). Also, actin inhibitors have been shown to block *E. invadens* encystation (22,23). So the role of actin in *Entamoeba* cyst chitin wall deposition was also investigated.

## Results and discussion

### Chitin wall formation starts from a single point and then spread over the whole cyst surface

*In vitro* encystation of *E. invadens* is an asynchronous process and takes nearly 48 to 72 hours to form a mature tetranucleated cyst. The progression of encystation is usually estimated by calculating the percentage of the detergent-resistant cysts. Cysts become detergent resistance only when the cyst wall formation is complete. The wall forming early stages are not resistant to detergent as staining with calcofluor white (CFW), the chitin-binding dye showed that the numbers of detergent-resistant cells were less than the total number of CFW positive encysting cells, especially in the early hours of encystations (Fig 1A). In order to identify chitin positive but detergent sensitive cells, encystation culture was examined at different time points (0 to 48 hr) by staining the chitin wall with CFW (Fig 1B). No fluorescence was observed on the encysting cells till the 9^th^ hour, but many cells with intense CFW fluorescence spots or cells with fluorescence covering only a portion of their surface were observed between 12 to 16^th^ hour’s encystation culture, indicating the initiation of partial cyst wall formation. By 24 to 48^th^ hours, the fluorescence was found all over the cell surface. These cells with incomplete chitin walls (Fig 1C) could be the intermediates in cyst wall formation. The numbers of such cells were counted manually using microscopy. The percentage of trophozoites (T), intermediate cyst with the partial wall (IC), and the cells with the complete or nearly complete wall (CC) are given in Fig 1D. Up to 20 hours of encystation, intermediate cysts (IC) comprised nearly 20% of the total cells and gradually decreased to about 5% at 24^th^ hour. These observations confirm that the chitin deposition started after 9 hours and actively took place between 12 to 24 hours of encystations. All the Chitin biosynthesis pathway genes and the genes of all other known protein components of the cyst wall were also found to be over-expressed during the same interval of encystations (17, 18).

**Fig 1.**
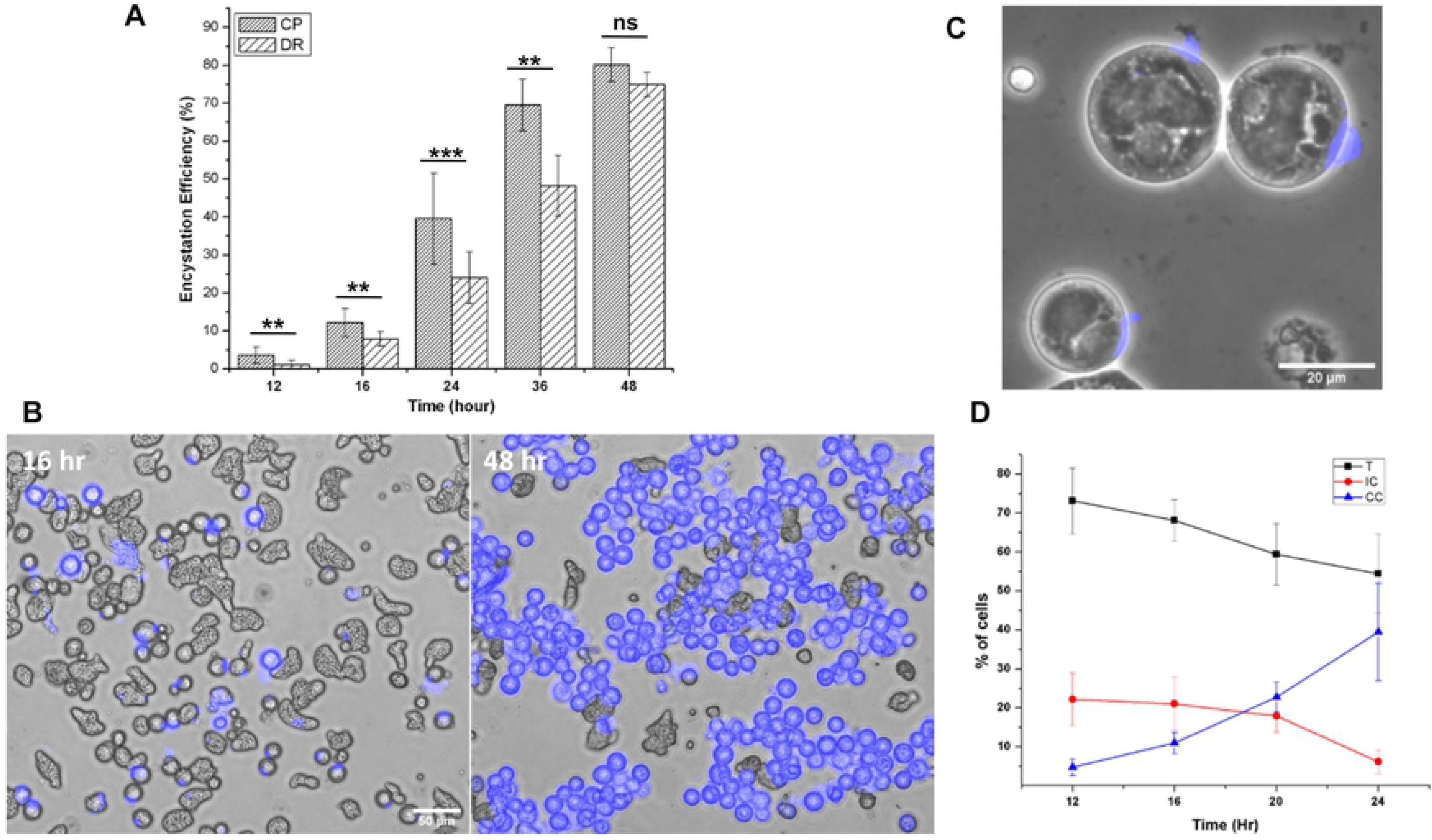
Cysts with incomplete chitin walls represent the intermediate stages in chitin wall formation. **(A)** During the early hours of encystation the number of detergent resistant cells was less than number of chitin positive cells even though both counting methods depended on the presence of chitin. **(B)** When treated with calcofluor white (CFW), early encystation culture (12, 16, and 20 hrs) showed cells with partial chitin wall and by 48th hours the chitin wall was complete (Scale bar: 50 μm). **(C)** CFW stained **e**ncysting cells showing partial chitin wall (Scale bar: 20 μm). **(D)** Percentage of trophozoites (T), Cysts with incomplete walls (IC) and complete walls (CC) at 12, 16, 20 and 24 hours. (Data are shown as mean ± SD for a minimum of 3 independent experiments. *p < 0.05, **p < 0.01, ***p < 0.001, ns not significant).

Comprehensive observation and analysis of CFW stained early encystation culture with confocal microscopy helped us to find a good number of possible intermediates of chitin wall formation. A probable sequence of wall formation is shown in Fig 2A. In most cells, wall formation was observed to start at one point, from which the chitin fibrils grew outwards and covered the whole surface. Similar cell wall formation was also observed in regenerating protoplasts of tobacco (24) and *Schizosaccharomyces pombe* (25,26). This pattern of chitin deposition might be due to the movement of wall synthesizing machinery at the site of wall formation, resembling the movement of cellulose synthase complex in plants (27) or maybe due to the self-assembly of wall components from a single point. Also, in a few encysting cells, the chitin wall seemed to start from multiple points and then spread all over the surface, finally overlapping and forming a continuous cyst wall (Fig 2B) (14).

**Fig 2.**
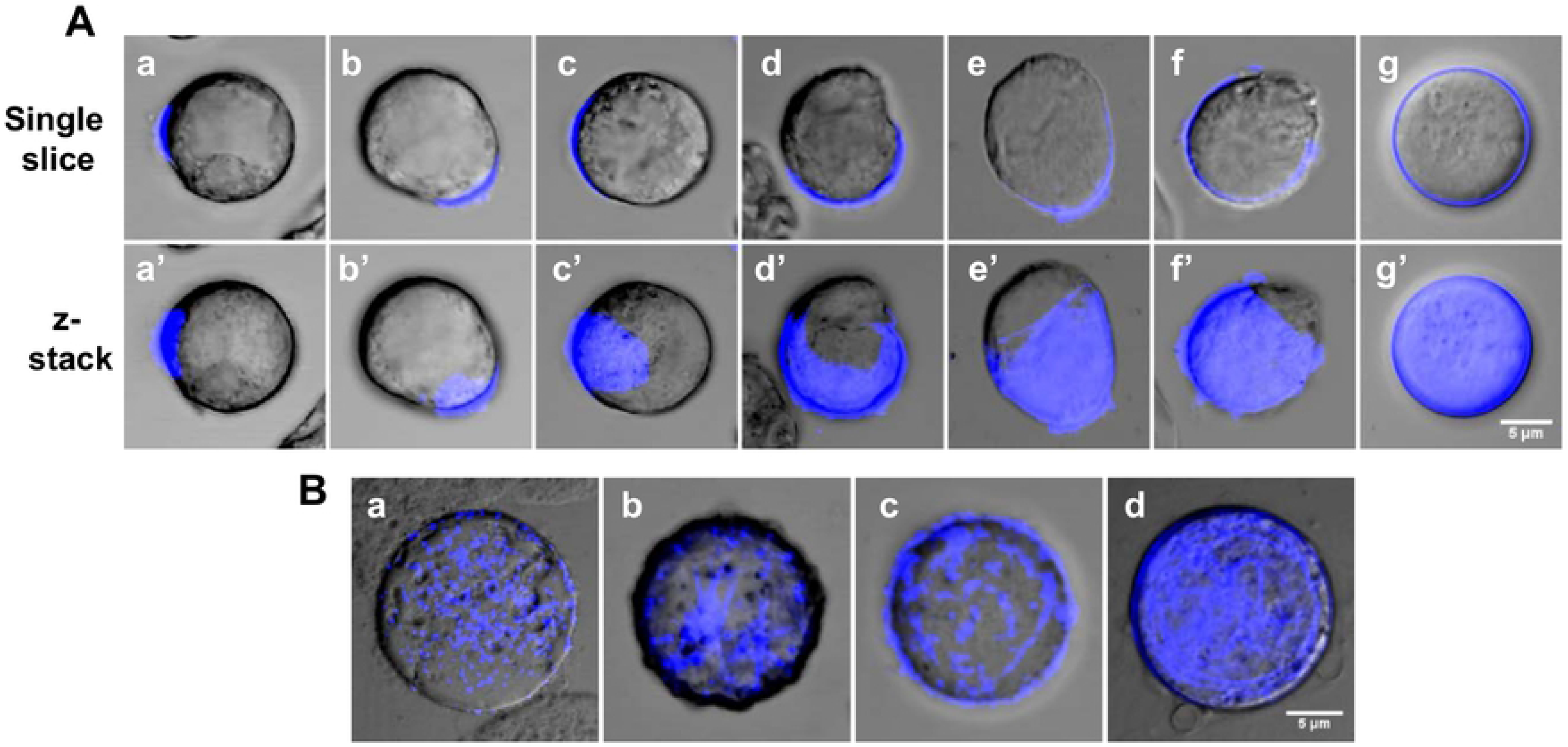
Possible sequence of chitin wall deposition. **(A)** All possible intermediates of chitin wall formation obtained from 9-16 hours of encystation observed with confocal microscopy. Upper panel (a to g) shows single slice and lower panel (a’ to g’) shows z stack. **(B)** In a few cells chitin formation started at multiple points (a) and their size increased with time (b, c), finally forming linear arrays on the cell surface (d). Scale bar: 5 μm.

Apart from the chitin wall, other important characteristics of a mature cyst are the formation of four nuclei and the chromatoid body (Fig 3A). To find out how these characteristics appear in the cysts with respect to the wall formation, all the three were simultaneously stained (Fig 3B). The chromatoid body and the nuclei were specifically stained with acridine orange and DAPI, respectively, and Alexafluor 633 tagged wheat germ agglutinin was used to stain the chitin wall. The chromatoid body was the first feature to appear in encysting cells; it appeared as multiple small green fluorescing structures in the cytoplasm by 6 hours and with time coalesced with each other to form a compact structure. Chitin wall deposition took place between 12 to 20 hours and was completed by the 24^th^ hour during which the chromatoid body acquired its final form. By the 48^th^ hour, the nuclear division has been completed to produce the tetranucleated mature cyst.

**Fig 3.**
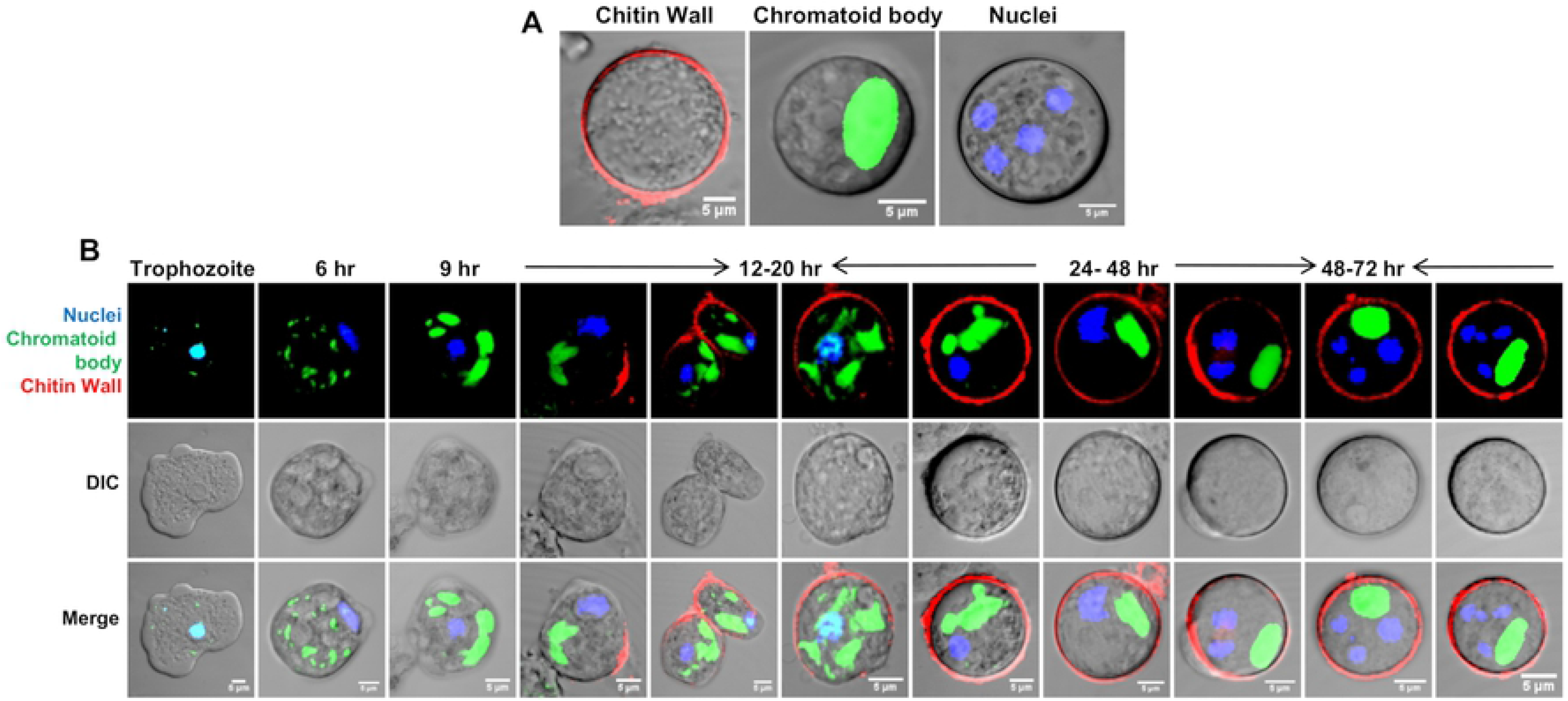
Chitin wall deposition with respect to nuclear division and chromatoid body formation. **(A)** The features of a mature *Entamoeba* cyst; chitin wall, chromatoid body and nucleus shown by staining with WGA, acridine orange and DAPI respectively. **(B)** Simultaneous observation of chromatoid body, nucleus and chitin wall in encysting cells showed that Chromatoid body appeared by 6^th^ hour as small aggregates which then coalesce with each other with time. Chitin wall deposition occurs between 12 to 20 hours and by the completion of chitin wall, chromatoid body was also fully formed. Chitin wall and chromatoid body were completed by 24 hour. Nuclear division occurs after the completion of chitin wall and produces the mature tetranucleated cyst. Scale bar: 5 μm.

## Localization of the chitin synthase on cyst surface guides chitin deposition

*Entamoeba invadens* contain two chitin synthase with different properties, and expression of both chitin synthase genes increased during the *in vitro* encystation (28). In the present work, we cloned and expressed truncated, chitin synthase 1 (EiCHS1), to raise polyclonal antibodies, which were then used for immunofluorescence microscopy (Fig 4 A). EiCHS1 was not observed in the trophozoite stage, but by 9^th^ hour EiCHS1 was localized in the cell distributed throughout the cytoplasm (Fig 4 Aa). As the encystations progressed, by 12-16 hours, EiCHS1 was found to be concentrated at one site (Fig 4 Ab), which was found to be the starting point for chitin deposition (Fig 4 Ac). Aggregation tendency of chitin synthase for proper functioning has already been reported elsewhere (29). Chitin synthases isolated from the tobacco hornworm midgut and mollusks were found to form a high molecular mass oligomeric complex (30,31). The cellulose synthase of plants formed a ‘‘rosette’’ complex to synthesis cellulose microfibrils (27). In *Entamoeba,* similar interactions between multiple chitin synthases on the membrane may be required for the secretion of chitin microfibrils. With time, more and more chitin synthase reached the plasma membrane and added chitin to the preexisting chitin wall and extending it and finally covering the entire surface (Fig 4 A d-g). In cysts with complete walls obtained from 24 hours of encystation (Fig 4 Ah) and in mature cysts from 48-72 hours (Fig 4 Ai), EiCHS1 co-localized with the chitin wall. The surface of the *Entamoeba* cyst usually contained an array of liner chitin fibrils, which can be seen from CFW staining (Fig 4 B). This patterned chitin deposition on the cyst surface was reflected in the immunolocalization of chitin synthase (Fig 4 B, C). These observations show that the chitin wall deposition may be guided by chitin synthase localization.

**Fig 4.**
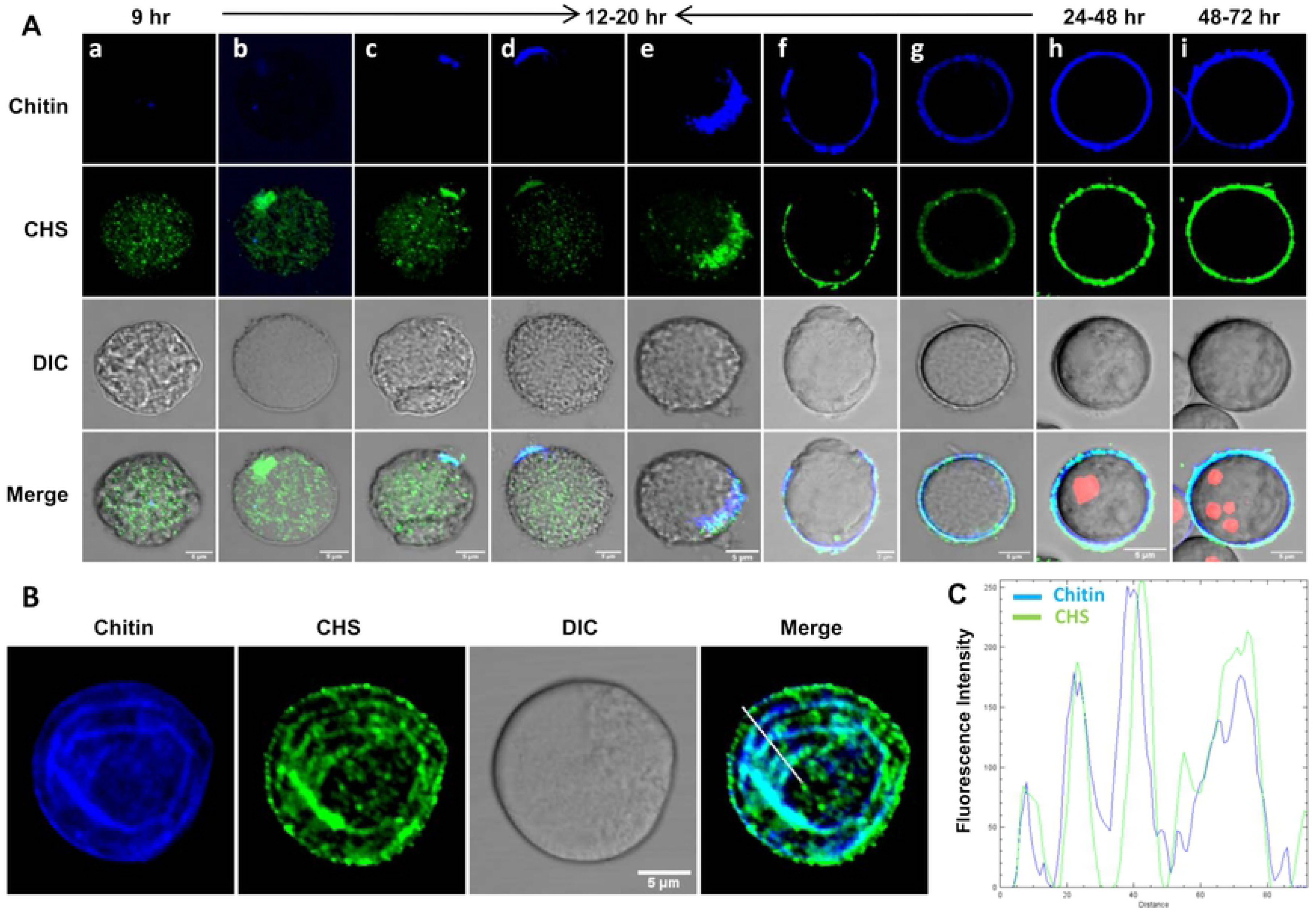
Immunolocalization of chitin synthase during encystation. **(A)** Chitin synthase was observed in the cytoplasm around 9^th^ hour of encysting cells. At 12^th^ hour onward it was observed to accumulate at certain points of plasma membrane and followed by chitin deposition at the same location. This incomplete wall then grew around the cell as chitin synthase added more chitin to it. In completed chitin wall of 24^th^ hour cysts, chitin synthase co-localized with chitin. In mature cysts, it also remained embedded in the wall. **(B)** On the cyst surface, chitin staining by CFW showed the presence of linear arrays of chitin fibrils. Localization of EiCHS 1 matched these patterns which can be seen from the plot of intensity values along the line drawn across the image **(C)**. Thus EiCHS 1 localization determines chitin deposition. Scale bar: 5 μm.

EiCHS 1 synthesized the chitin fibrils at the cell surface, and once the wall was completed, it remained associated with the chitin wall. Such localization of the chitin synthases in the wall itself has also been reported in fungi like *Mucor rouxii* and *Ustilago maydis* (32,33). Whether the chitin synthases are secreted to the wall or remained physically trapped in the secreted chitin chains (34) is not clear. Thus, like the lectin components, chitin synthase also becomes part of the wall instead of remaining in the plasma membrane.

Previously it has been shown that during the encystation, chitin was made in intracellular secretory vesicles and then deposited on the cyst wall (14,35). In this study, we have demonstrated that in the cyst, the chitin is synthesized by membrane-bound chitin synthase enzyme. Recently, a very similar model for cellulose microfibril synthesis by membrane-bound cellulose synthase has been proposed in *Achanthamoeba* cyst (36). We believe both the processes are taking place simultaneously in *Entamoeba.* To get the complete picture of chitin synthesis and deposition, the role of chitin synthase 2 also needs to be explored.

### Revisiting the Wattle and daub model of the cyst wall (Chitin, chitinase, and Jessie lectin assemble into the cyst wall from one starting point on the foundation made of Jacob lectin)

In this study, we have revisited cyst wall formation and noticed the same sequence of events during cyst wall formation as refer to Wattle and daub model (14). Here, we followed encystation by co-localizing known cyst wall proteins like Jacob, Jessie, and chitinase using their respective antibodies (14,37,38) with the growing chitin wall. Co-localizations of Jacob, chitinase, and Jessie independently with chitin at different time points of encystations are demonstrated in Fig 5, Fig 6A, and 6B, respectively. This study indicates that the complete, as well as the incomplete wall, were in accordance with the Wattle and daub model with respect to cyst wall composition, sequences of events, and organization of the components.

**Fig 5.**
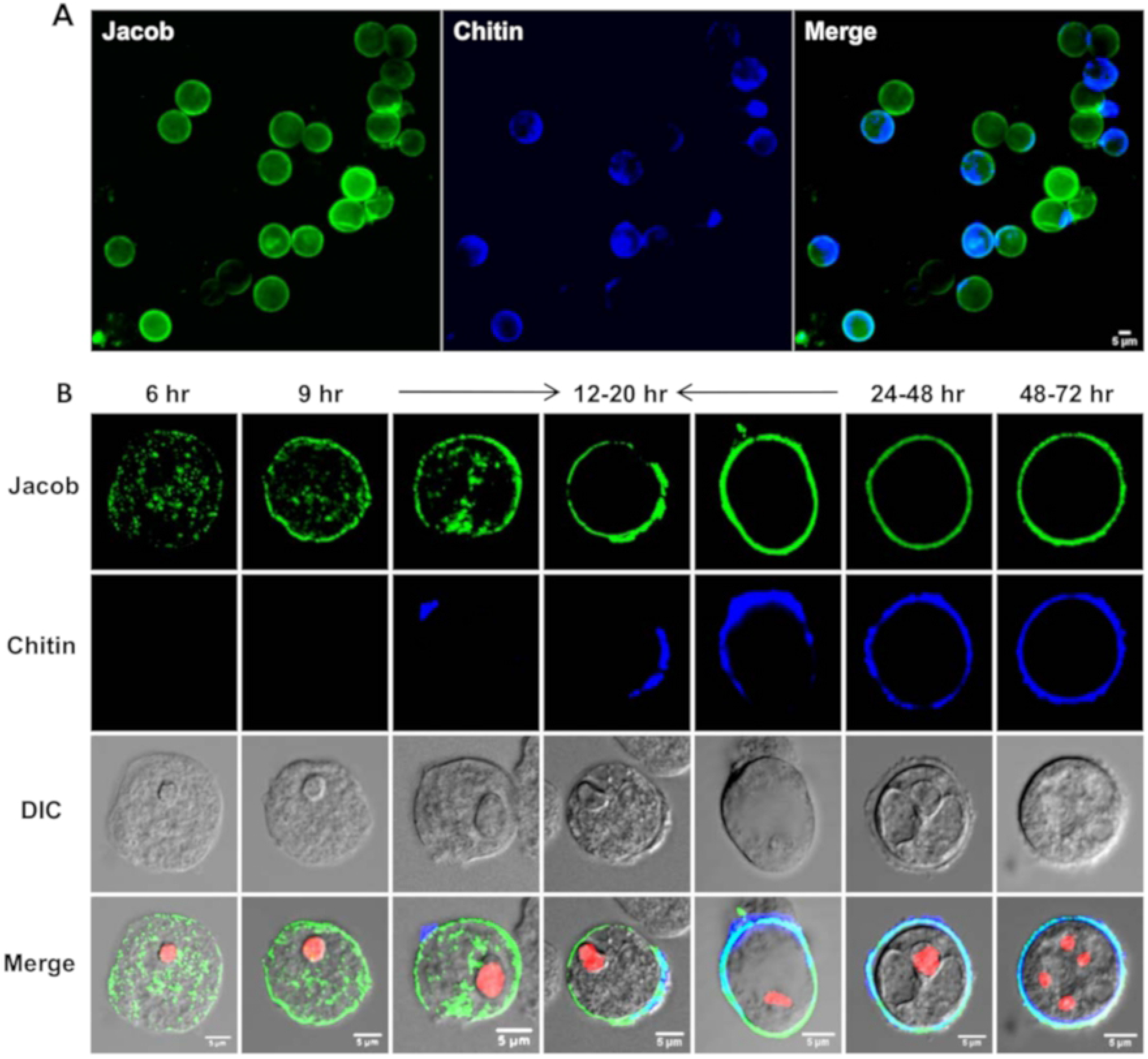
Immunolocalization of Jacob during encystation. **(A)**12 hour encystation culture showing presence of Jacob lectin on the surface of most cells with the chitin being deposited on the cell surface starting from one point. **(B)** Jacob lectin was observed in the cytoplasm by 6^th^ hour and by 9^th^ hour it was found on the membrane. Between 12 to 20 hours chitin wall was deposited on this Jacob layer, starting from one point. In 24^th^ hour and 48^th^ hour mature cyst, Jacob co-localized with the chitin cyst wall as it became a part of the wall. Scale bar: 5 μm.

**Fig 6.**
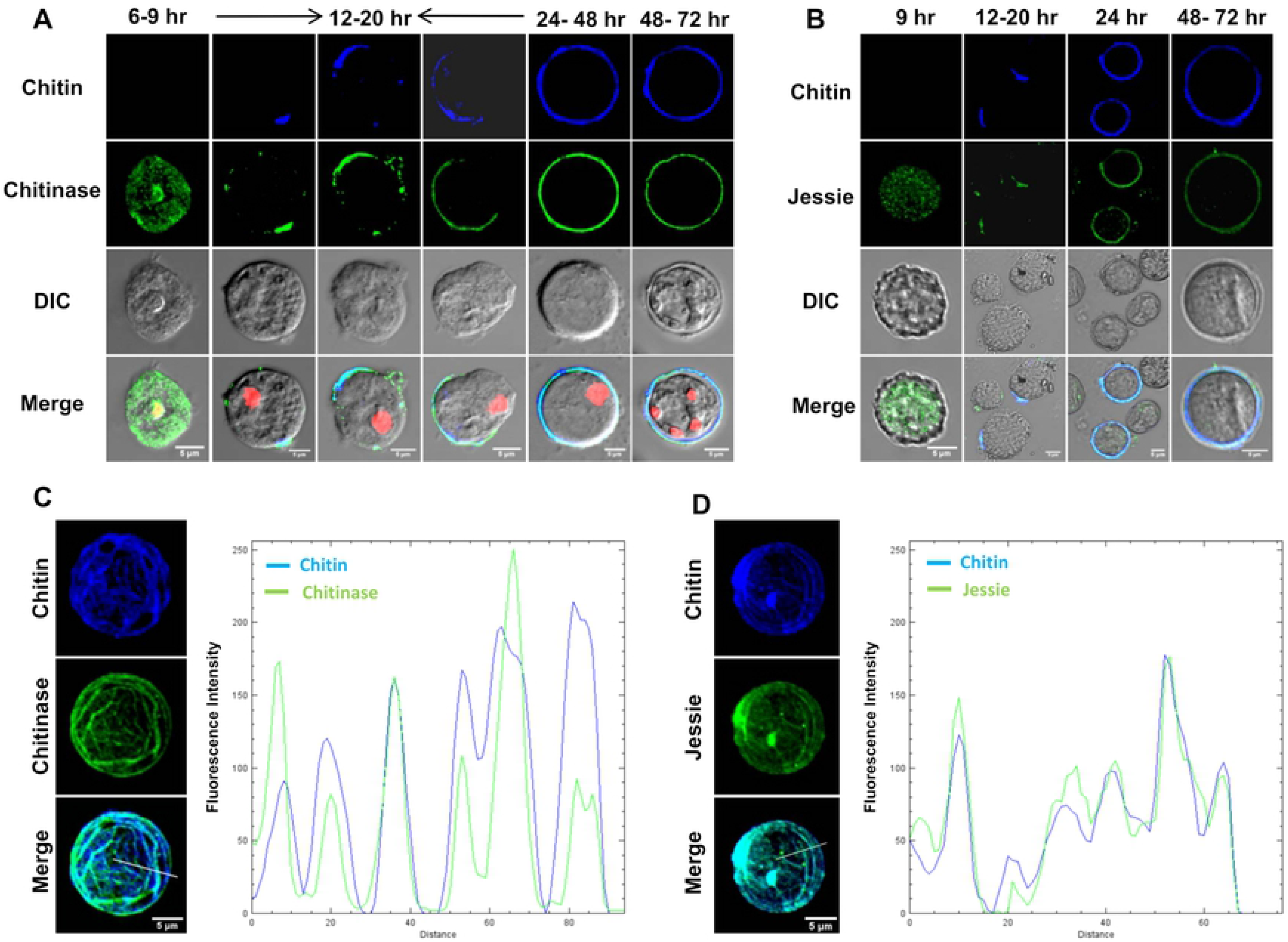
Immunolocalization of Chitinase and Jessie during encystation. **(A)** Chitinase lectin was observed in the cytoplasm by 6^th^ hour. Chitinase always co-localized with the chitin in cysts with incomplete walls obtained from 12-20 hours. After wall was completed, it remained embedded in the wall. **(B)** Jessie was found to appear in the vesicle of cytoplasm of encysting cells at around 9 hour of encystations and was also found in the incomplete wall from 12-20 hours of encystation culture. It was also found in the cysts with complete walls from 24^th^ hour and in the mature cyst from 48-72 hour encystation. **(C, D)** Deposition pattern of chitinase **(C)** and Jessie (**D)** on the cyst surface followed the pattern of chitin deposition, as observed by CFW staining. The plots of intensity values along the line drawn across the confocal images also show the co-localization of chitin with the lectins. Scale bar: 5 μm.

In contrary to the previously described model, we have noticed that Jessie expresses at about 9^th^ hour of encystation and also found to co-localize with the growing chitin wall (Fig 6B). The early expression of Jessie also is in consensus with the expression profile of different cyst wall proteins, including Jessie (17, 18). However, the expression of Jessie is relatively low and hard to detect unless it is co-localized with chitin. We believe that in all the previous reports of cyst wall biosynthesis, partial cyst wall formation was overlooked, and so the expression of Jessie during the early hour of encystations. This is the first report where all the cyst wall proteins (CWP) were co-localized with cyst wall chitin, and that helped us to spot the CWP clearly.

Our observations clearly indicate that the Jacob and chitin synthase deposition on the cell surface are chitin independent, whereas the deposition of chitinase or Jessie is chitin dependent. Encysting cells with ridge-like chitin all over the surface, along with co-localized chitinase and Jessie, were also visible (Fig 6 C, D). Those woolen balls like structure are also reported by Chatterjee et al. (2007). Those cells are most likely originated by multipoint initiation of cyst wall formation and also much brighter with respect to cells with partially formed walls under the fluorescence microscope. The population of these brighter cells is much more attractive to be imaged, and so the less bright populations were ignored in previous reports.

Encystation took place only inside the multicellular aggregates found in the encystation culture, and it could be possible that the components of the cyst wall are secreted into the intercellular space where they underwent self-assembly into complete chitin wall on the cell surface from a starting point (Fig 7). Jessie lectin has a unique C-terminal domain that appears to promote self-aggregation (14). Also, the ionic attraction between negatively charged Jacob, Jessie, and chitinase and positively charged chitosan could be mediating such self-assembly. This may also explain the importance of cell aggregation in the cyst formation, as within the aggregate, the concentration of these secreted cell wall components would be high, promoting the wall assembly (39).

**Fig 7.**
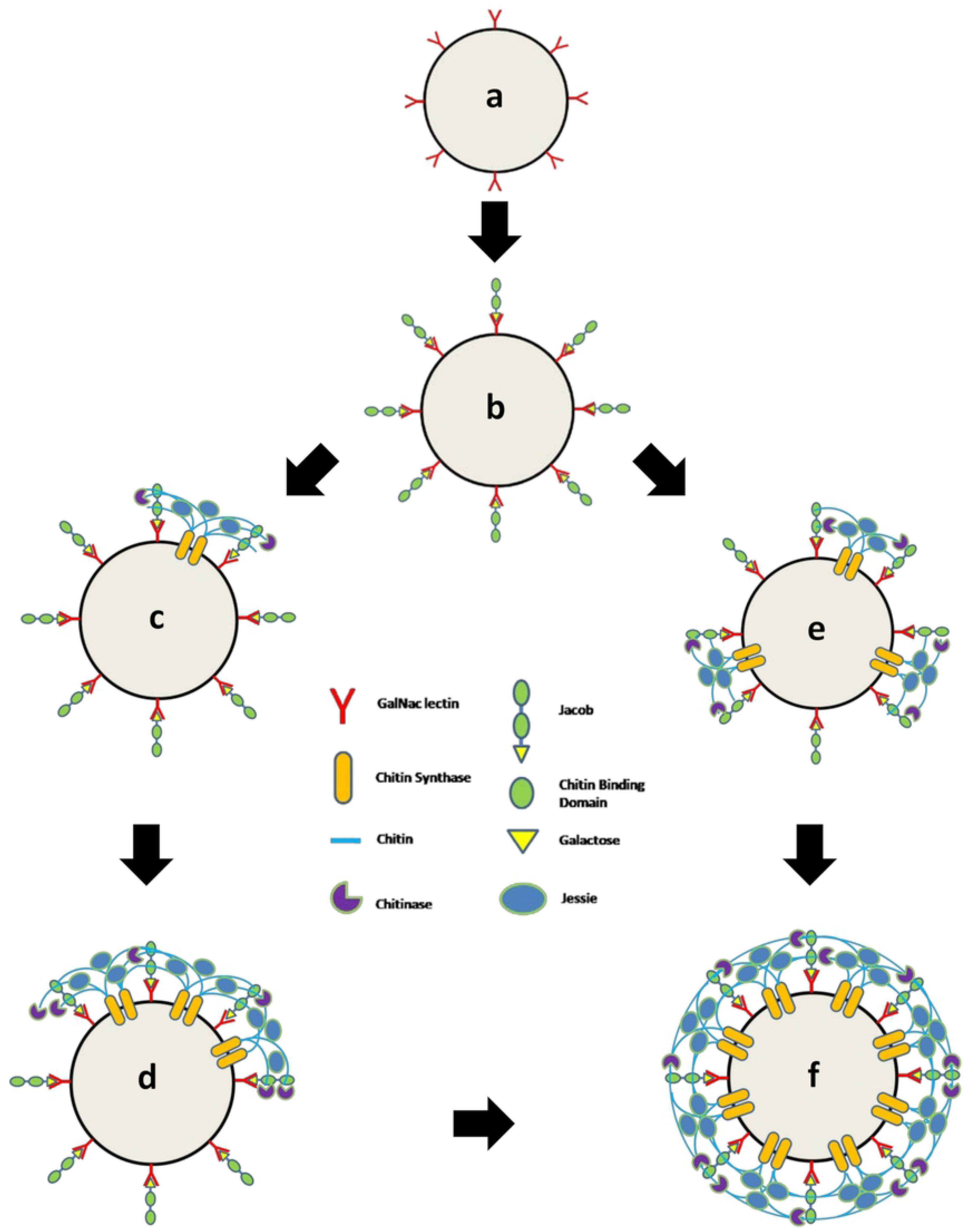
Model of chitin wall formation. Jacob lectin is secreted and binds to the Gal/GalNAc lectins (a, b). Chitin synthase produce the chitin fibrils at one point (c) or multiple points (e). Secreted chitinases and Jessie binds to this framework. Cyst wall grows from these points (d) and finally covers the whole surface (f). The major pathway is a-b-c-d-f and minor pathway is a-b-e-f.

### During encystation, F-actin was reorganized into the cortical region where it remained until the completion of the chitin wall

In the trophozoite stage of *Entamoeba*, the dynamic cytoskeleton is involved in motility, pseudopodia formation, and phagocytosis. During the encystation, *E. invadens* undergo morphological changes from highly motile trophozoites to an immobile spherical cyst, which shows the involvement of the actin cytoskeleton in the stage conversion. *Entamoeba* lacks cytoplasmic microtubules, and so the cell morphology is determined by the actin cytoskeleton alone (21). Specific inhibitors of actin, like Cytochalasin D and jasplakinolide, have been shown to inhibit encystation of *E. invadens* (22,23). To find the involvement of actin in the cyst chitin wall formation, actin was localized using Rhodamine-phalloidin in encysting cells.

In the trophozoites, actin was localized mainly in the phagocytic/ pinocytic invaginations found all over the surface (Fig 8A). Within one hour of encystation, all the actin was observed to polymerize at the cortical region (Fig 8B). Uniform actin polymerization at the cortical region is reported to cause isotropic contraction during mitotic cell rounding (40), and the same may also be responsible for the round morphology of the cysts. Further studies indicated that between 12 to 24 hours, chitin synthase started to accumulate on the cortical actin that eventually guides the chitin synthesis and chitin wall formation over this cortical actin (Fig 8C). As shown previously (Fig 4A), more chitin synthase reached the cyst surface, depositing chitin and extending the wall. Thus the actin cytoskeleton may be acting as a scaffold to support the wall formation, similar to the cytoskeleton guided cellulose deposition in plant cell walls (41). The cortical actin cytoskeleton may also be involved in transport and secretion of cyst wall components, as observed during the encystation of *Giardia intestinalis* (42). Changes in the cortical actin localization were observed only after the completion of the wall. When cysts obtained from 24^th^ and 48 hours were stained with Rhodamine phalloidin (Fig. 9 A, B), actin remained in the cell cortex in a few cysts, but in most cases, it formed aggregates of different sizes in the cytoplasm. Most of the encysting cells demonstrated that cortical actin gets disassembled after the wall formation was completed, indicating a temporal relationship between cortical actin and chitin deposition.

**Fig 8.**
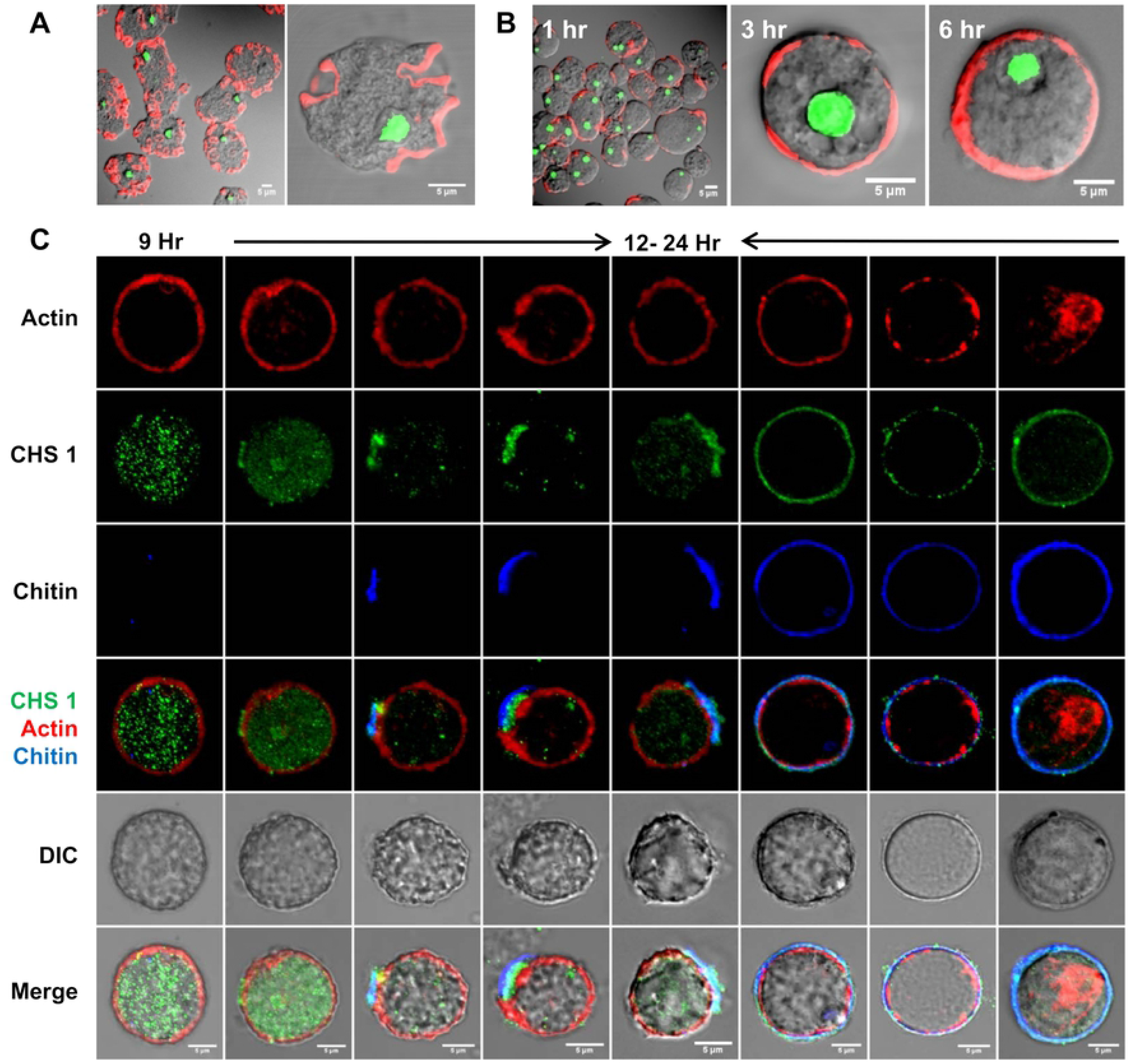
Reorganization of actin cytoskeleton during encystation. **(A)** In trophozoites actin was found mainly in the phagocytic and pinocytic invaginations. **(B)** Within one hour of encystation, all the actin polymerized at the cortical region. **(C)** Chitin synthase vesicles are found in the cytoplasm by 9^th^ hour. Between 12^th^ to 24 hours chitin synthase was observed to accumulate on the cortical actin and where chitin synthesis started. The chitin wall then grows as more chitin syntheses migrate towards cell membrane and synthesized chitin. Chitin synthase remained as the wall component after the completion of the wall, but the cortical actin was observed to disassemble afterwards. Scale bar: 5 μm.

**Fig 9.**
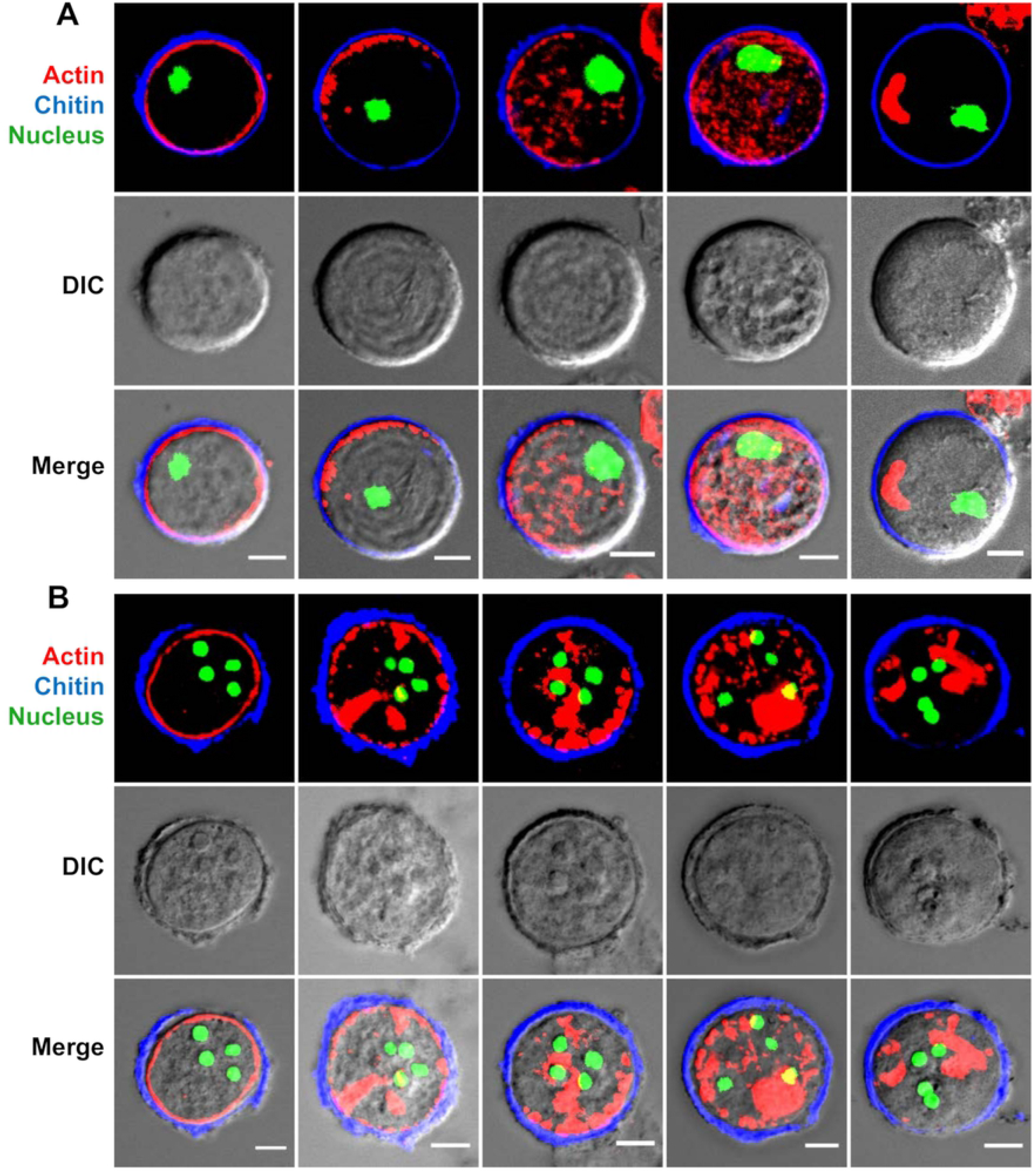
Localization of cortical actin after wall formation. Conversion of cortical actin ring to actin aggregates in the cytoplasm after the completion of wall **(A)** in cysts at 24 hours and **(B)** in 48-72 hour mature cysts. Scale bar: 5 μm.

### 2, 3-Butanedione monoxime treatment inhibited cortical actin formation and produced wall-less cysts and cysts with defective walls

To understand the role of cortical actin in *Entamoeba* cyst wall deposition, actin polymerization was disrupted by 2,3-Butanedione monoxime (BDM). BDM has been used to study the synthesis and organization of yeast chitin cell wall (43,44) and it has been observed to disorganize polar distribution of actin patches in *Schizosaccharomyces pombe* resulting in an uneven cell wall (43). BDM inhibits actomyosin contractility by blocking the ATPase activity of myosin II (45) and also found to specifically inhibit the Myosin-dependent functions in *E. histolytica* (46).

At 20 mM concentration, BDM inhibited the rearrangement of cellular actin into the cell cortex, as shown by Rhodamine phalloidin staining. In BDM treated cells, actin was observed to have formed an aggregate in the cytoplasm (Fig 10A). When added to the encystation culture, BDM dose-dependently inhibited the encystation, with 20 mM BDM reducing the encystation efficiency to < 5% (Fig 10B). In the encystation culture, the cysts are formed only inside the galactose ligand-mediated cell aggregates formed during encystation, and cell signaling required for encystation takes place in these aggregates (47). The addition of BDM did not inhibit the cell aggregates, but only a few chitin walled cysts were found in these aggregates, as shown by CFW staining (Fig 10C). However, in BDM treated cells, no change in the expression level of cyst wall proteins was observed by RT-PCR (Fig 10D) and CFW staining showed many cells with disordered and patchy localization of chitin on their surface (Fig 10E). Further staining of BDM treated cells for the nucleus, and chromatoid body revealed that most cells contained the chromatoid body and four nuclei indicating the cells entered the encystation process and formed mature cysts (Fig 10F). The presence of many tetranucleate cells with the chromatoid body indicated that BDM treatment affected cyst wall formation without preventing the cyst maturation. The presence of cysts with aberrant walls and ‘‘wall-less cysts’’ in BDM treated cultures showed that the cortical actin cytoskeleton is necessary for proper chitin deposition. These observations also showed that chitin wall deposition and other developmental events took place independently of each other.

**Fig 10.**
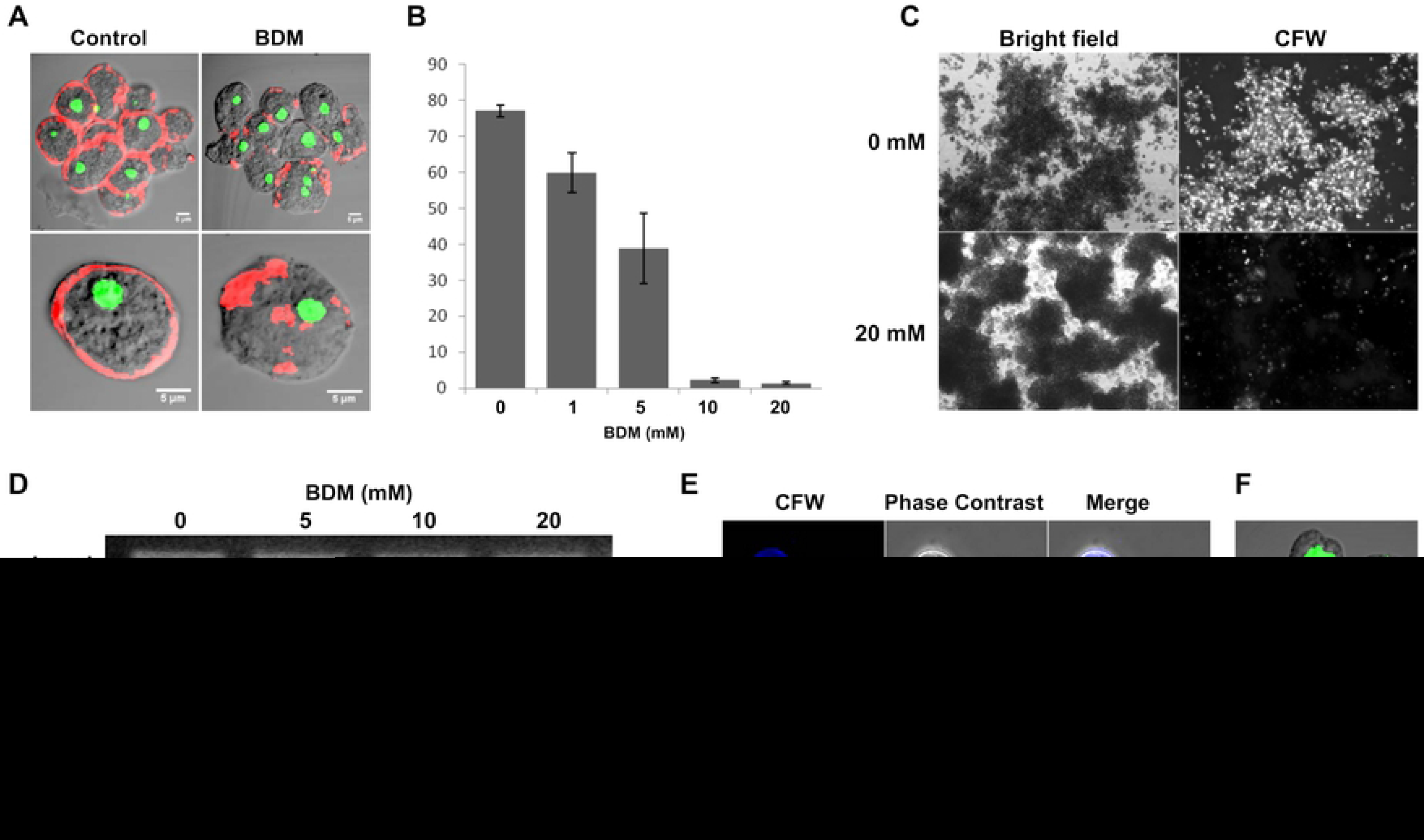
Effects of 2, 3-Butanedione monoxime on encystation. **(A)** Cortical actin was found in encysting cells by 1 hour, but in BDM treated cells actin was found as cytoplasmic aggregates. (Scale bar: 5 μm). **(B)** BDM dose dependently reduced encystation efficiency. **(C)** Cysts were formed only inside the cell aggregates during encystation. In BDM treated culture, the aggregates still formed but fewer cysts were found inside the cell aggregates. Scale bar: 100 μm. **(D)** Semi quantitative RT-PCR with mRNA isolated from 24^th^ hour encysting cells showed that all cyst wall genes are expressed in BDM treated encystation culture. **(E)** Control and BDM treated cysts obtained from 48^th^ hour encystation culture after staining with CFW. BDM treatment caused aberrant chitin deposition. **(F)** Nuclear staining with PI and indirect staining of chromatoid body with FITC showed large number of tetranucleated “wall less cysts” in culture. Scale bar: 10 μm. ( Data are shown as mean ± SD for a minimum of 3 independent experiments.)

### Inhibition of cortical actin formation affected both localization and activity of chitin synthase

BDM treated encystation culture contained both “wall-less cysts” and cysts with abnormal walls. Instead of forming a uniform chitin wall (Fig 11A a), in BDM treated cells, chitin was deposited in an irregular fashion (Fig 11A b), but in most cases, it was found to be accumulated on one side (Fig 11A c, d). To find how the cortical actin disruption affects the cyst wall formation, the cyst wall proteins were localized (Fig 11B). Immunostaining of Jacob in BDM treated cells showed no change in its localization, it was present on the cyst surface in both ‘‘wall-less cysts’’ and cyst with aberrant walls, just like in the normal cysts (Fig 11B, Jacob). Jacob secretion or it’s binding to the Gal/GalNAc lectin, and thus the ‘‘foundation’’ phase of encystation was not affected by BDM. Immunolocalization of chitinase and Jessie in BDM treated cells with their respective antibodies showed that these proteins co-localized with the aberrant chitin walls like the normal chitin walls (Fig 11B, Chitinase, Jessie). This could be because these proteins were secreted to the extracellular space and get attached to the chitin fibrils through their chitin-binding domains. Thus the secretions of cell wall components were not affected, but only the deposition of chitin, which indicated a link between cortical actin and chitin synthase localization. In temperature-sensitive actin mutants of yeasts, dysfunction of actin led to the formation of an aberrant wall by delocalizing the chitin synthase (48,49). In *Neurospora crassa,* actin inhibitors caused mislocalization of chitin synthase, indicating the role of actin in traffic and localization of chitin synthase (50).

**Fig 11.**
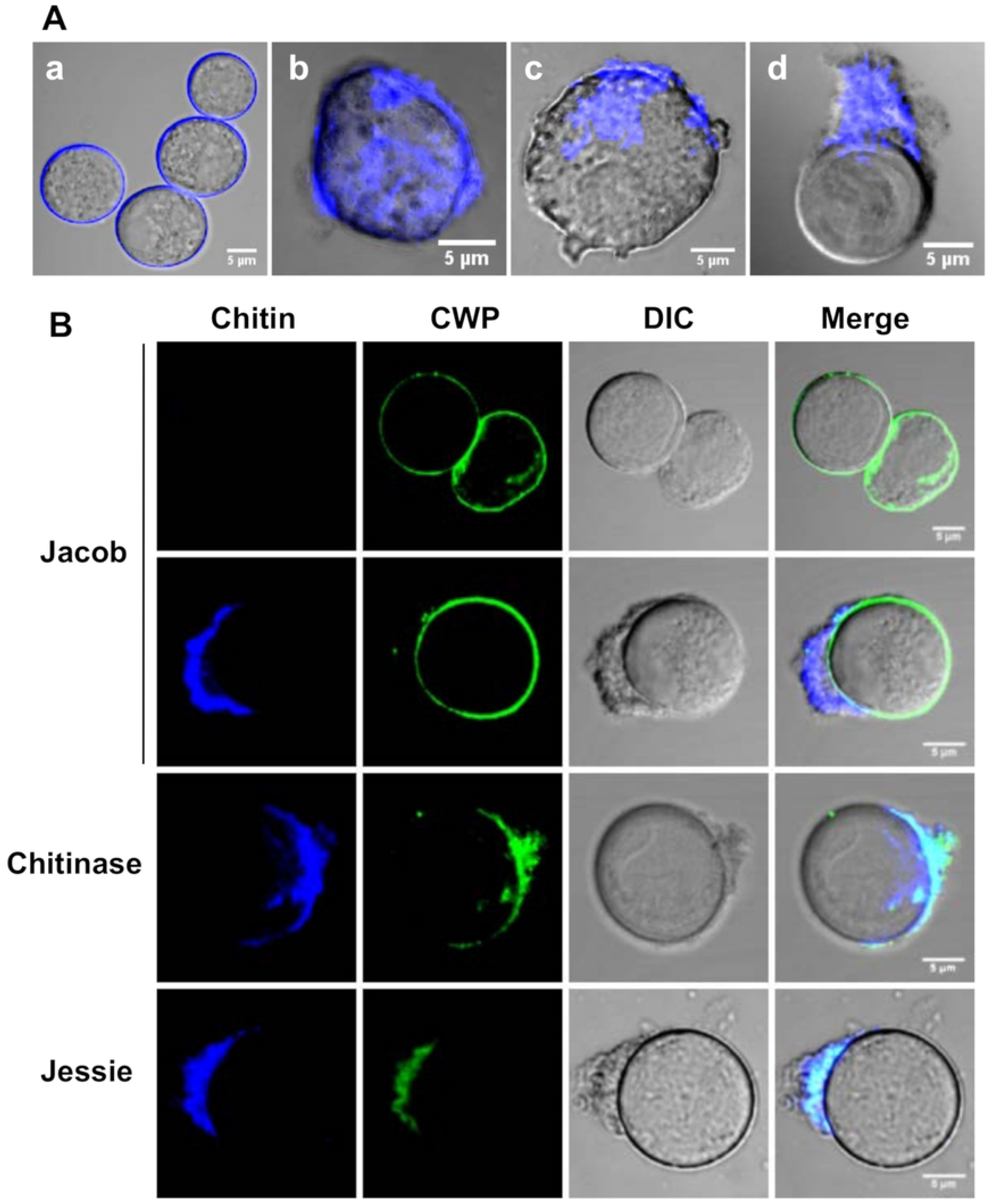
Localization of chitin and cyst wall proteins (CWP) in BDM treated cysts. **(A)** chitin wall is of uniform thickness in control cysts, (a). BDM treatment caused aberrant cyst wall formation and uneven chitin deposition (b) and clustering of chitin fibrils to one end of cell (c, d). **(B)** Confocal microscopy showed that in BDM treated cyst no change in the secretion or localization of Jacob. Jacob was found on the surface of cysts with no chitin wall or with aberrant chitin deposition. Chitinase and Jessie were always co localized with chitin since they were secreted outside and contained chitin binding domains. Scale bar: 5 μm.

Immunolocalization of chitin synthase (EiCHS1) in BDM treated cells obtained from 72-hour encystation culture showed that it was present on the surface of most cells, both cysts with aberrant chitin walls (Fig 12 A, B) and the wall-less cysts (Fig 12 C, D). Thus the effects of BDM on cell wall deposition were resulting from its inhibition of chitin synthase localization and/or its activity, indicating a relationship between the cortical actin cytoskeleton and chitin synthase 1. In *Aspergillus nidulans,* chitin synthase was concentrated at the sites of chitin synthesis like hyphal tips and septation sites. This localization required interaction between the actin cytoskeleton and myosin motor-like domain (MMD) of chitin synthase, and the MMD-actin interaction was necessary for both localization and function of chitin synthase (51). The function of the cortical actin cytoskeleton may be to anchor chitin synthases and act as a scaffold to maintain the structure. Since EiCHS 1 did not contain any MMD, further studies are required to find the exact nature of the interaction between the actin cytoskeleton and chitin synthase, if there is any.

**Fig 12.**
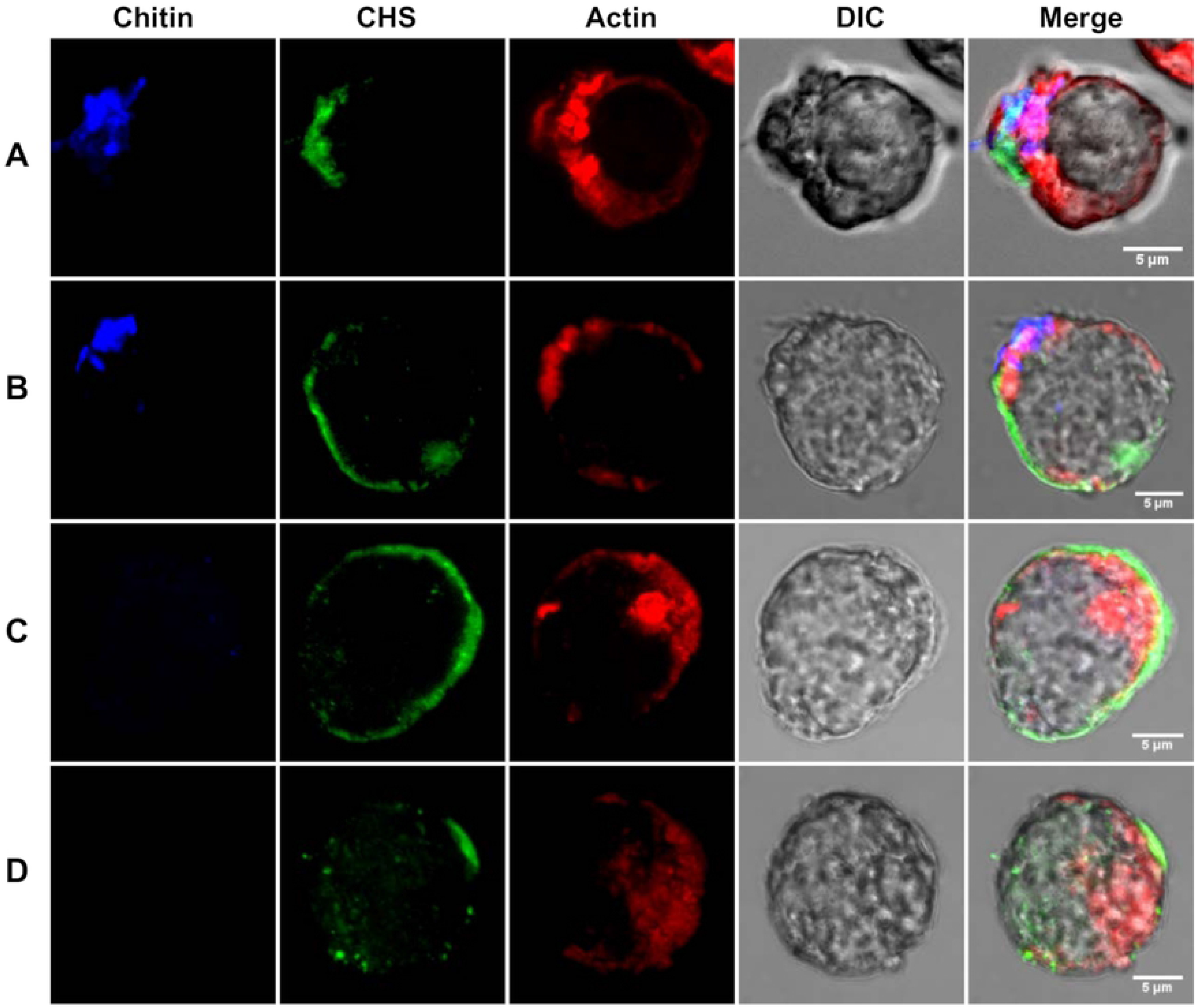
Immunolocalization of Chitin synthase in BDM treated cells. Actin remained delocalized in BDM treated cells. Uneven Chitin deposition was observed in some cells **(A, B)**; others did not form chitin even though chitin synthase was observed on the cell surface (**C, D**). Scale bar: 5 μm.

In conclusion, this study explored the cyst wall formation in more detail in relation to chitin deposition on cyst wall and chromatoid body formation, and the role of Chitin synthase has been reported for the first time. Various aspects of actin scaffold with respect to cyst wall formation have also been demonstrated. Wattle and daub model of *Entamoeba* cyst wall biosynthesis has been revised, and the modified model is explained in Fig 7.

### Materials and methods

### Cells and reagents

*Entamoeba invadens* strain IP-1 was maintained in TYI-S-33 medium containing 10% adult bovine serum (HiMedia) and 3% Diamond vitamin mix at 25°C. DAPI, Propidium iodide, calcofluor white, Alexafluor 633 tagged wheat germ agglutinin, and acridine orange were purchased from Sigma-Aldrich. 2,3-Butanedione monoxime was purchased from HiMedia. TRITC conjugated phalloidin was purchased from Molecular Probes, Invitrogen, USA. Antibodies against Chitinase and Chitin synthase were synthesized in the lab. Jacob and Jessei3 antibodies were kindly provided by Dr. John Samuelson.

### Chitin synthase antibody preparation

To generate antibody for localization of chitin synthase 1 (EiCHS1), gene fragment encoding a part of catalytic domain in EiCHS1 was cloned and heterologously expressed in bacterial cell as a protein of 17.5kD. Primers 5’GGATCCCACTCGCACGAAATCTTCTTT3’ and 5’CTCGAGGTACCCTGTATCCCACCCAG 3’ were used to amplify a region from 921bp-1374bp of Eichs1 gene and the amplified product was cloned into pET-21a(+) vector. The construct was transformed into C43 (DE3) bacterial strains for protein expression. The polyclonal antibody against this expressed protein was commercially raised in rabbit by ABENEX Pvt. Ltd. India.

### Encystation

To prepare the encystation (LG 47), TYI medium without glucose was prepared and diluted to 2.12 times and then completed with 5% heat inactivated adult bovine serum, 1.5 % vitamin mix and antibiotics, penicillin and streptomycin. Mid log phase trophozoites were chilled on ice for 10 minutes to detach the cells from the culture tube wall and harvested by centrifugation at 500 ×g for 5 min at 4° C. These cells were washed 3 times with LG 47 and then cell number was adjusted to 5 × 10^5^ trophozoites per ml and incubated at 25 °C.

### Estimation of Encystation efficiency

Encystation efficiency was calculated in two different ways, one based on detergent resistance and other based the number of chitin positive cells after staining with fluorescent chitin binding dyes. To estimate the encystation efficiency from detergent resistance, the cells were harvested from the encystation culture and was counted using haemocytometer. It was treated with 0.1% Sarkosyl for 10 minutes to remove trophozoites and then counted the number of detergent-resistant cysts from which the encystation efficiency was calculated. To measure the percentage of cysts with incomplete and complete chitin wells, CFW stained (30 μM) encystation culture was taken on a slide and images were taken at random under phase contrast and UV light. From these images, the percentage of fluorescing cysts, both incomplete and complete were calculated. To observe the formation of cysts inside the aggregates, 30 μM CFW was directly added to the encystation medium, incubated for 10 minutes, and then observed under fluorescence microscope. Each encystation experiment was repeated a minimum of three times, and at least 60 cells were counted to calculate encystation efficiency at each time point.

### Cell staining

To stain the chitin wall, the cells were treated with calcofluor white (CFW) at 30 μM concentration for 5 minutes before fixing the cell as the dead cells were observed to stain non-specifically. The chitin wall was also stained with Alexafluor 633-conjugated wheat germ agglutinin (WGA) at a concentration of 20 μg per ml. After PBS wash to remove the CFW, cells were fixed with 4% (w/v) paraformaldehyde in PBS for 10 minutes and then permeabilized in 0.5% (v/v) Triton X-100 in PBS for 5 minutes. DAPI (0.5 μg/ ml), Sytox Green (1 μM) and PI (10 μg/ ml) were used to stain the nucleus. Acridine Orange at 10 μM concentration and Fluorescein isothiocyanate (by non-specific binding) were used to stain the chromatoid body. For actin localization, permeabilized cells were blocked with 2% (w/v) BSA and stained with TRITC conjugated phalloidin (Molecular Probes, Invitrogen, USA) according to manufacturer’s protocols. To localize the cyst wall proteins CFW stained, fixed and permeabilized cells were, blocked with 2% (w/v) BSA and then incubated with appropriate dilution of cell wall protein antibody followed by Fluorescein isothiocyanate (FITC) -conjugated goat anti-rabbit secondary antibody and examined by fluorescence and confocal microscopy.

### Microscopy

Olympus IX51 inverted light microscope with camera attachment and photo-editing software (Image Pro Discovery) and Olympus FV1000 confocal microscope with Fluoview software was used for fluorescence imaging. Image analysis was then done using ImageJ software (NIH). To find the co-localization of chitin fibrils with cyst wall lectins and chitin synthase, RGB Profiler plugin was used.

### RNA Isolation and RT-PCR

Total RNA was isolated from different hours of encystation culture using TRIzol reagent (Invitrogen). After treating total RNA with DNaseI (Takara), first-strand cDNA was synthesized using MuLV super RT reverse transcriptase (BioBharati) according to the manufacturer’s specifications. Using first-strand cDNA as template, semi-quantitative RT-PCR was done to find the expression of chitin wall proteins using Premix Taq (Xceleris) DNA polymerase and the following primer pairs:

#### Primers for RT-PCR

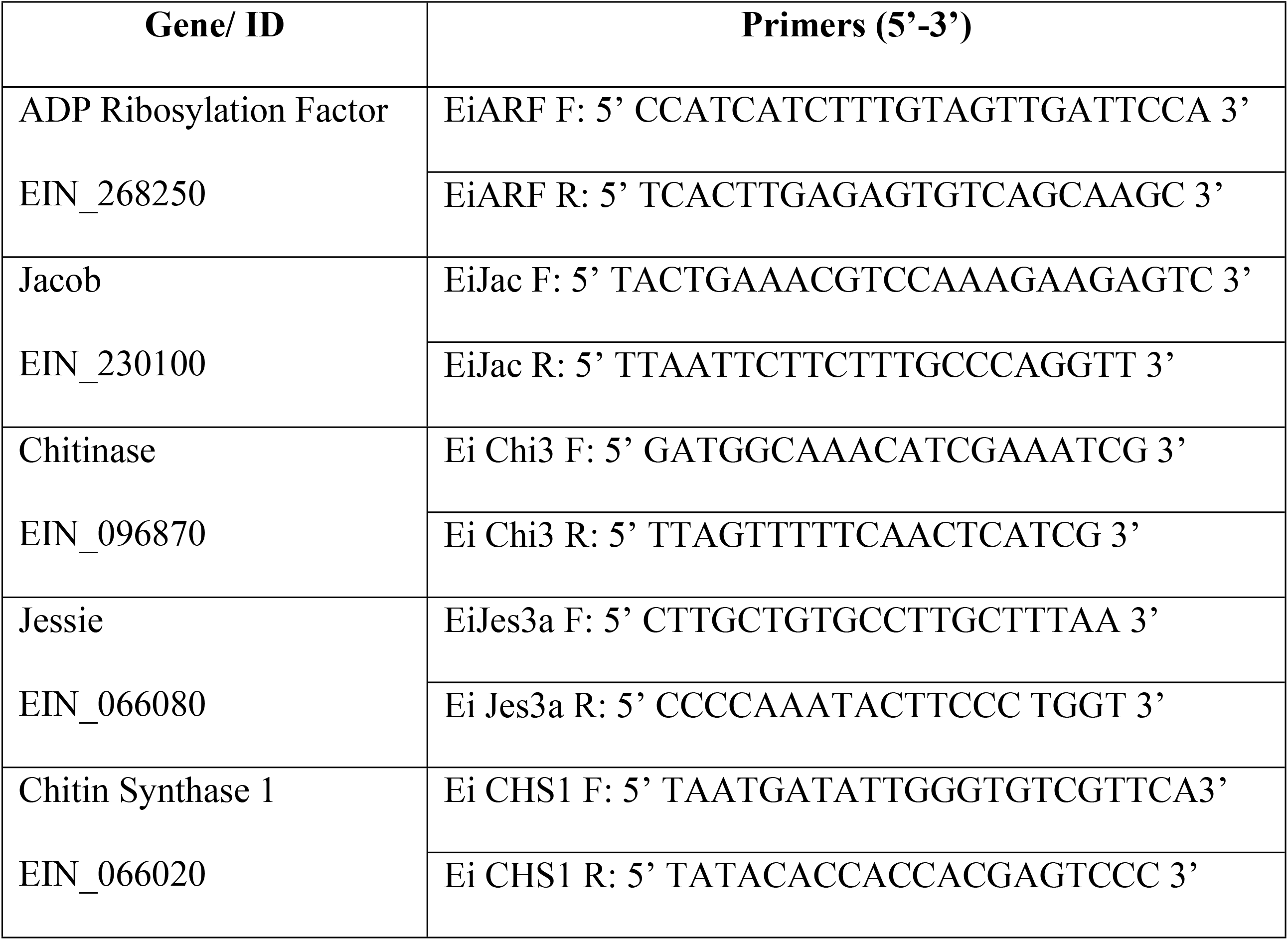

The PCR products were analyzed by running in a 1.5% agarose gel. EiARF was used as an internal control.

## Acknowledgments

This work was partially funded by Ministry of Human Resources, and Ministry of Health and Family Welfare, Govt. of India. DK is recipient of Junior Research Fellowship from Indian Council of Medical Research, India. SN is recipient of Junior Research fellowship from UGC. Anti Jacob and anti Jessie 3 antibodies were kindly provided by Dr. John Samuelson, Boston University Medical Center, Boston, USA. Authors thank FIST, DST, Government of India for confocal facility. The authors declare no conflict of interest.

## Author Contributions

DK and SKG designed research; DK carried out the encystation, confocal microscopy, immunolocalization and BDM treatment studies; SN cloned and expressed the chitin synthase fragment and generated polyclonal antibody; DK and SKG wrote the paper.

## References

1. WHO/PAHO/UNESCO report. A consultation with experts on amoebiasis. Mexico City, Mexico 28-29 January, 1997. Epidemiol Bull. 1997 Mar;18(1):13–4. PMID: 9197085

2. Wassmann C, Hellberg A, Tannich E, Bruchhaus I. Metronidazole resistance in the protozoan parasite *Entamoeba histolytica* is associated with increased expression of iron-containing superoxide dismutase and peroxiredoxin and decreased expression of ferredoxin 1 and flavin reductase. J Biol Chem. 1999 Sep 10;274(37):26051–6. doi: 10.1074/jbc.274.37.26051. PMID: 10473552

3. Orozco E, Gómez C, Pérez DG. Physiology and molecular genetics of multidrug resistance in *Entamoeba histolytica*. Drug Resist Updat. 1999 Jun;2(3):188–97. doi:10.1054/drup.1999.0087. PMID: 11504490

4. Larocque R, Nakagaki K, Lee P, Abdul-Wahid A, Faubert GM. Oral immunization of BALB/c mice with *Giardia duodenalis* recombinant cyst wall protein inhibits shedding of cysts. Infect Immun. 2003 Oct;71(10):5662–9. PMID: 14500486 doi: 10.1128/iai.71.10.5662-5669.2003

5. Lee P, Faubert GM. Expression of the *Giardia lamblia* cyst wall protein 2 in Lactococcus lactis. Microbiology. 2006 Jul 1;152(7):1981–90. PMID: 16804173 doi: 10.1099/mic.0.28877-0

6. Feng X-M, Zheng W-Y, Zhang H-M, Shi W-Y, Li Y, Cui B-J, et al. Vaccination with bivalent DNA vaccine of α1-Giardin and CWP2 delivered by attenuated *Salmonella typhimurium*reduces trophozoites and cysts in the feces of mice infected with *Giardia lamblia*. PLoS One. 2016 Jun 22;11(6):e0157872. PMID: 27332547 doi: 10.1371/journal.pone.0157872

7. Sanchez L, Enea V, Eichinger D. Identification of a developmentally regulated transcript expressed during encystation of Entamoeba invadens. Mol Biochem Parasitol. 1994 Sep;67(1):125–35. doi: 10.1016/0166-6851(94)90102-3 PMID: 7838173

8. Chia M-Y, Jeng C-R, Hsiao S-H, Lee A-H, Chen C-Y, Pang VF. *Entamoeba invadens* Myositis in a Common Water Monitor Lizard *(Varanus salvator)*. Vet Pathol. 2009 Jul 9;46(4):673–6. doi: 10.1354/vp.08-VP-0224-P-CRPMID: 19276058

9. Wang Z, Samuelson J, Clark CG, Eichinger D, Paul J, Van Dellen K, et al. Gene discovery in the *Entamoeba invadens* genome. Mol Biochem Parasitol. 2003 Jun;129(1):23–31. doi: 10.1016/s0166-6851(03)00073-2PMID: 12798503

10. Ehrenkaufer GM, Haque R, Hackney JA, Eichinger DJ, Singh U. Identification of developmentally regulated genes in *Entamoeba histolytica*: insights into mechanisms of stage conversion in a protozoan parasite. Cell Microbiol. 2007 Jun;9(6):1426–44. PMID: 17250591 doi: 10.1111/j.1462-5822.2006.00882.x

11. Ghosh SK, Van Dellen KL, Chatterjee A, Dey T, Haque R, Robbins PW, et al. The Jacob2 Lectin of the *Entamoeba histolytica*cyst wall binds chitin and is polymorphic. PLoS Negl Trop Dis. 2010 Jul 20;4(7):e750. doi: 10.1371/journal.pntd.0000750PMID: 20652032

12. Arroyo-Begovich A, Cárabez-Trejo A, Ruíz-Herrera J. Identification of the structural component in the cyst wall of *Entamoeba invadens*. J Parasitol. 1980 Oct;66(5):735–41. PMID: 7463242

13. Van Dellen KL, Chatterjee A, Ratner DM, Magnelli PE, Cipollo JF, Steffen M, et al. Unique posttranslational modifications of chitin-binding lectins of *Entamoeba invadens* cyst walls. Eukaryot Cell. 2006 May;5(5):836–48. doi: 10.1128/EC.5.5.836-848.2006PMID: 16682461

14. Chatterjee A, Ghosh SK, Jang K, Bullitt E, Moore L, Robbins PW, et al. Evidence for a “Wattle and Daub” Model of the cyst wall of *Entamoeba*. PLoS Pathog. 2009 Jul 3;5(7):e1000498. PMID: 19578434. doi: 10.1371/journal.ppat.1000498

15. Nayak S, Ghosh SK. Nucleotide sugar transporters of *Entamoeba histolytica* and *Entamoeba invadens* involved in chitin synthesis. Mol Biochem Parasitol. 2019 Dec;234:111224. doi.org/10.1016/j.molbiopara.2019.111224

16. Das S, Van Dellen K, Bulik D, Magnelli P, Cui J, Head J, et al. The cyst wall of *Entamoeba invadens* contains chitosan (deacetylated chitin). Mol Biochem Parasitol. 2006 Jul;148(1):86–92. doi: 10.1016/j.molbiopara.2006.03.002PMID: 16621070

17. De Cádiz AE, Jeelani G, Nakada-Tsukui K, Caler E, Nozaki T. Transcriptome analysis of encystation in *Entamoeba invadens.* Bogyo M, editor. PLoS One. 2013 Sep 11;8(9):e74840. PMID: 24040350. doi: 10.1371/journal.pone.0074840.

18. Ehrenkaufer GM, Weedall GD, Williams D, Lorenzi HA, Caler E, Hall N, et al. The genome and transcriptome of the enteric parasite *Entamoeba invadens*, a model for encystation. Genome Biol. 2013;14(7):R77.PMID: 23889909 doi: 10.1186/gb-2013-14-7-r77

19. Harris SD, Morrell JL, Hamer JE. Identification and characterization of *Aspergillus nidulans* mutants defective in cytokinesis. Genetics. 1994 Feb;136(2):517–32.PMID: 8150280

20. Gabriel M, Kopecka M, Svoboda A. Structural rearrangement of the actin cytoskeleton in regenerating protoplasts of budding yeasts. J Gen Microbiol. 1992 Oct 1;138(10):2229–34. doi: 10.1099/00221287-138-10-2229

21. Vayssié L, Vargas M, Weber C, Guillén N. Double-stranded RNA mediates homology-dependant gene silencing of γ-tubulin in the human parasite *Entamoeba histolytica*. Mol Biochem Parasitol. 2004 Nov 1;138(1):21–8. PMID: 15500912 doi: 10.1016/j.molbiopara.2004.07.005

22. Makioka A, Kumagai M, Ohtomo H, Kobayashi S, Takeuchi T. Effect of cytochalasin D on the growth, encystation, and multinucleation of *Entamoeba invadens*. Parasitol Res. 2000 Jun 23;86(7):599–602. doi.org/10.1007/PL00008536

23. Makioka A, Kumagai M, Ohtomo H, Kobayashi S, Takeuchi T. Effect of jasplakinolide on the growth, encystation, and actin cytoskeleton of *Entamoeba histolytica* and *Entamoeba invadens*. J Parasitol. 2001 Apr;87(2):399–405. PMID: 11318572 doi: 10.1645/0022-3395(2001)087[0399:EOJOTG]2.0.CO;2

24. Hahne G, Herth W, Hoffmann F. Wall formation and cell division in fluorescence-labelled plant protoplasts. Protoplasma. 1983 Jun;115(2–3):217–21.doi.org/10.1007/BF01279812

25. Osumi M, Sato M, Ishijima SA, Konomi M, Takagi T, Yaguchi H. Dynamics of cell wall formation in fission yeast,*Schizosaccharomyces pombe*. Fungal Genet Biol. 1998 Jun;24(1–2):178–206.MID: 9742201 doi: 10.1006/fgbi.1998.1067

26. Kobori H, Yamada N, Taki A, Osumi M. Actin is associated with the formation of the cell wall in reverting protoplasts of the fission yeast *Schizosaccharomyces pombe*. J Cell Sci. 1989 Dec;94(Pt 4):635–46. PMID: 2630560

27. Kimura S, Laosinchai W, Itoh T, Cui X, Linder C, Brown R, et al. Immunogold labeling of rosette terminal cellulose-synthesizing complexes in the vascular plant *Vigna angularis*. Plant Cell. 1999 Nov;11(11):2075–86. PMID: 10559435 doi: 10.1105/tpc.11.11.2075

28. Campos-Góngora E, Ebert F, Willhoeft U, Said-Fernández S, Tannich E. Characterization of chitin synthases from *Entamoeba*. Protist. 2004 Sep;155(3):323–30.PMID: 15552059 doi: 10.1078/1434461041844204

29. Ruiz-Herrera J, Bartnicki-Garcia S. Synthesis of cell wall microfibrils in vitro by a “soluble” chitin synthetase from *Mucor rouxii*. Science (80). 1974 Oct 25;186(4161):357–9. PMID: 1058485 doi: 10.1073/pnas.72.7.2706

30. Maue L, Meissner D, Merzendorfer H. Purification of an active, oligomeric chitin synthase complex from the midgut of the tobacco hornworm. Insect Biochem Mol Biol. 2009 Sep;39(9):654–9. PMID: 19576988 doi: 10.1016/j.ibmb.2009.06.005

31. Weiss IM, Lüke F, Eichner N, Guth C, Clausen-Schaumann H. On the function of chitin synthase extracellular domains in biomineralization. J Struct Biol. 2013 Aug 1;183(2):216–25. PMID: 23643908 doi: 10.1016/j.jsb.2013.04.011

32. McMurrough I, Flores-Carreon A, Bartnicki-Garcia S. Pathway of chitin synthesis and cellular localization of chitin synthetase in *Mocor rouxii*. J Biol Chem. 1971 Jun 25;246(12):3999–4007.PMID: 5561471

33. Ruiz-Herrera J, Xoconostle-Cázares B, Reynaga-Peña CG, León-Ramírez C, Cárabez-Trejo A. Immunolocalization of chitin synthases in the phytopathogenic dimorphic fungus *Ustilago maydis*. FEMS Yeast Research. 2006. p. 999–1009. PMID: 17042749 doi: 10.1111/j.1567-1364.2006.00133.x

34. Kang MS, Elango N, Mattia E, Au-Young J, Robbins PW, Cabib E. Isolation of chitin synthetase from *Saccharomyces cerevisiae*. Purification of an enzyme by entrapment in the reaction product. J Biol Chem. 1984 Dec 10;259(23):14966–72. PMID: 6238967

35. Chávez-Munguía B, Cristóbal-Ramos AR, González-Robles A, Tsutsumi V, Martínez-Palomo A. Ultrastructural study of *Entamoeba invadens* encystation and excystation. J Submicrosc Cytol Pathol. 2003 Jul;35(3):235–43. PMID: 14690171

36. Garajová M, Mrva M, Vaškovicová N, Martinka M, Melicherová J, Valigurová A. Cellulose fibrils formation and organisation of cytoskeleton during encystment are essential for *Acanthamoeba* cyst wall architecture. Sci Rep. 2019 Dec 1;9(1):1–21. doi: 10.1038/s41598-019-41084-6

37. Ghosh SK, Field J, Frisardi M, Rosenthal B, Mai Z, Rogers R, et al. Chitinase secretion by encysting *Entamoeba invadens* and transfected *Entamoeba histolytica* trophozoites: Localization of secretory vesicles, endoplasmic reticulum, and Golgi apparatus. Infect Immun. 1999 Jun;67(6):3073–81. PMID: 10338523

38. Dey T, Basu R, Ghosh SK. *Entamoeba invadens*: cloning and molecular characterization of chitinases. Exp Parasitol. 2009 Nov;123(3):244–9. doi: 10.1016/j.exppara.2009.07.008.

39. Frederick JR, Petri WA. Roles for the galactose-/N-acetylgalactosamine-binding lectin of *Entamoeba* in parasite virulence and differentiation. Glycobiology. 2005 Dec 1;15(12):53R–59R. PMID: 16037494 doi: 10.1093/glycob/cwj007

40. Stewart MP, Helenius J, Toyoda Y, Ramanathan SP, Muller DJ, Hyman AA. Hydrostatic pressure and the actomyosin cortex drive mitotic cell rounding. Nature. 2011 Jan 2;469(7329):226–30. doi: 10.1038/nature09642

41. Paredez AR, Somerville CR, Ehrhardt DW. Visualization of cellulose synthase demonstrates functional association with microtubules. Science. 2006 Jun 9; 312(5779):1491–5. PMID: 16627697 doi: 10.1126/science.1126551

42. Paredez AR, Assaf ZJ, Sept D, Timofejeva L, Dawson SC, Wang C-JR, et al. An actin cytoskeleton with evolutionarily conserved functions in the absence of canonical actin-binding proteins. Proc Natl Acad Sci. 2011 Apr 12;108(15):6151–6.PMID: 21444821 doi: 10.1073/pnas.1018593108

43. Steinberg G, McIntosh JR. Effects of the myosin inhibitor 2,3-butanedione monoxime on the physiology of fission yeast. Eur J Cell Biol. 1998 Dec;77(4):284–93.PMID: 9930653 doi: 10.1016/S0171-9335(98)80087-3

44. Rivera-Molina FE, González-Crespo S, Cruz YM-D la, Ortiz-Betancourt JM, Rodríguez-Medina JR. 2,3-Butanedione monoxime increases sensitivity to Nikkomycin Z in the budding yeast *Saccharomyces cerevisiae*. World J Microbiol Biotechnol. 2006 Mar 12;22(3):255–60.PMID: 25382940 doi: 10.1007/s11274-005-9028-x

45. Sambrano GR, Fraser I, Han H, Ni Y, O’Connell T, Yan Z, et al. Navigating the signalling network in mouse cardiac myocytes. Nature. 2002 Dec 12;420(6916):712–4. PMID: 12478303 doi: 10.1038/nature01306

46. Manning-Cela R, Marquez C, Franco E, Talamas-Rohana P, Meza I. BFA-sensitive and insensitive exocytic pathways in *Entamoeba histolytica* trophozoites: their relationship to pathogenesis. Cell Microbiol. 2003 Dec;5(12):921–32. PMID: 14641177

47. Coppi A, Merali S, Eichinger D. The enteric parasite *Entamoeba*uses an autocrine catecholamine system during differentiation into the infectious cyst Stage. J Biol Chem. 2002 Mar 8;277(10):8083–90. PMID: 11779874 doi: 10.1074/jbc.M111895200

48. Takagi T, Ishijima SA, Ochi H, Osumi M. Ultrastructure and behavior of actin cytoskeleton during cell wall formation in the fission yeast *Schizosaccharomyces pombe*. J Electron Microsc (Tokyo). 2003;52(2):161–74. PMID: 7773392 doi: 10.1099/13500872-141-4-891PMID: 12868587 doi: 10.1093/jmicro/52.2.161

49. Gabriel M, Kopecka M. Disruption of the actin cytoskeleton in budding yeast results in formation of an aberrant cell wall. Microbiology. 1995 Apr 1;141(4):891–9. PMID: 7773392 doi: 10.1099/13500872-141-4-891

50. Sánchez-León E, Verdín J, Freitag M, Roberson RW, Bartnicki-Garcia S, Riquelme M. Traffic of chitin synthase 1 (CHS-1) to the Spitzenkörper and developing septa in hyphae of *Neurospora crassa*: Actin dependence and evidence of distinct microvesicle populations. Eukaryot Cell. 2011 May;10(5):683–95. PMID: 21296914 doi: 10.1128/EC.00280-10

51. Takeshita N, Ohta A, Horiuchi H. CsmA, a class V chitin synthase with a myosin motor-like domain, is localized through direct interaction with the actin cytoskeleton in *Aspergillus nidulans*. Mol Biol Cell. 2005 Apr;16(4):1961–70. PMID: 15703213 doi: 10.1091/mbc.E04-09-0761

